# Differential Sensitivity of Midline Patterning to Mitosis during and after Primitive Streak Extension

**DOI:** 10.1101/2024.10.25.620280

**Authors:** Zhiling Zhao, Rieko Asai, Takashi Mikawa

## Abstract

**Background:** Midline establishment is a fundamental process during early embryogenesis for *Bilaterians*. Midline patterning in nonamniotes can occur without mitosis, through Planar Cell Polarity (PCP) signaling. By contrast, amniotes utilize both cell proliferation and PCP signaling for patterning early midline landmark, the primitive streak (PS). This study examined their roles for midline patterning at post PS-extension.

**Results:** In contrast to PS extension stages, embryos under mitotic arrest during the post PS-extension preserved notochord (NC) extension and Hensen’s node (HN)/PS regression judged by both morphology and marker genes, although they became shorter, and laterality was lost. Remarkably, no or background level of expression was detected for the majority of PCP core components in the NC-HN-PS area at post PS-extension stages, except for robustly detected *prickle-1*. Morpholino knockdown of Prickle-1 showed little influence on midline patterning, except for suppressed embryonic growth. Lastly, associated with mitotic arrest-induced size reduction, midline tissue cells displayed hypertrophy.

**Conclusion:** Thus, the study has identified at least two distinct mitosis sensitivity phases during early midline pattering: One is PS extension that requires both mitosis and PCP, and the other is mitotic arrest-resistant midline patterning with little influence by PCP at post PS-extension stages.

## Introduction

The bilateral body plan of *Bilaterian* critically depends on midline establishment. In non-amniote vertebrates, a sperm entry site predicts the future midline.^1,2^ Subsequent midline patterning, including axial mesoderm development, in part utilizing the PCP gene network,^3–7^ can take place even under mitotic arrest.^8^ Intercalation of mesodermal cells that results in convergent extension depends on non-canonical Wnt/PCP signaling.^7,9^

By contrast, midline development of amniote embryos requires both mitotic activities and PCP signaling,^10–14^ as exemplified in the formation of the PS,^15^ the first midline tissue formed as a conduit of amniote gastrulation^16^ (Figure 1**A**, Movie S1). Thus, midline patterning of amniote embryos consists of two distinct components; one is an evolutionarily conserved mechanism, such as utilizing PCP signaling, and the other is an evolutionarily new component, e.g., a requirement of mitotic activities.

**Figure 1.**
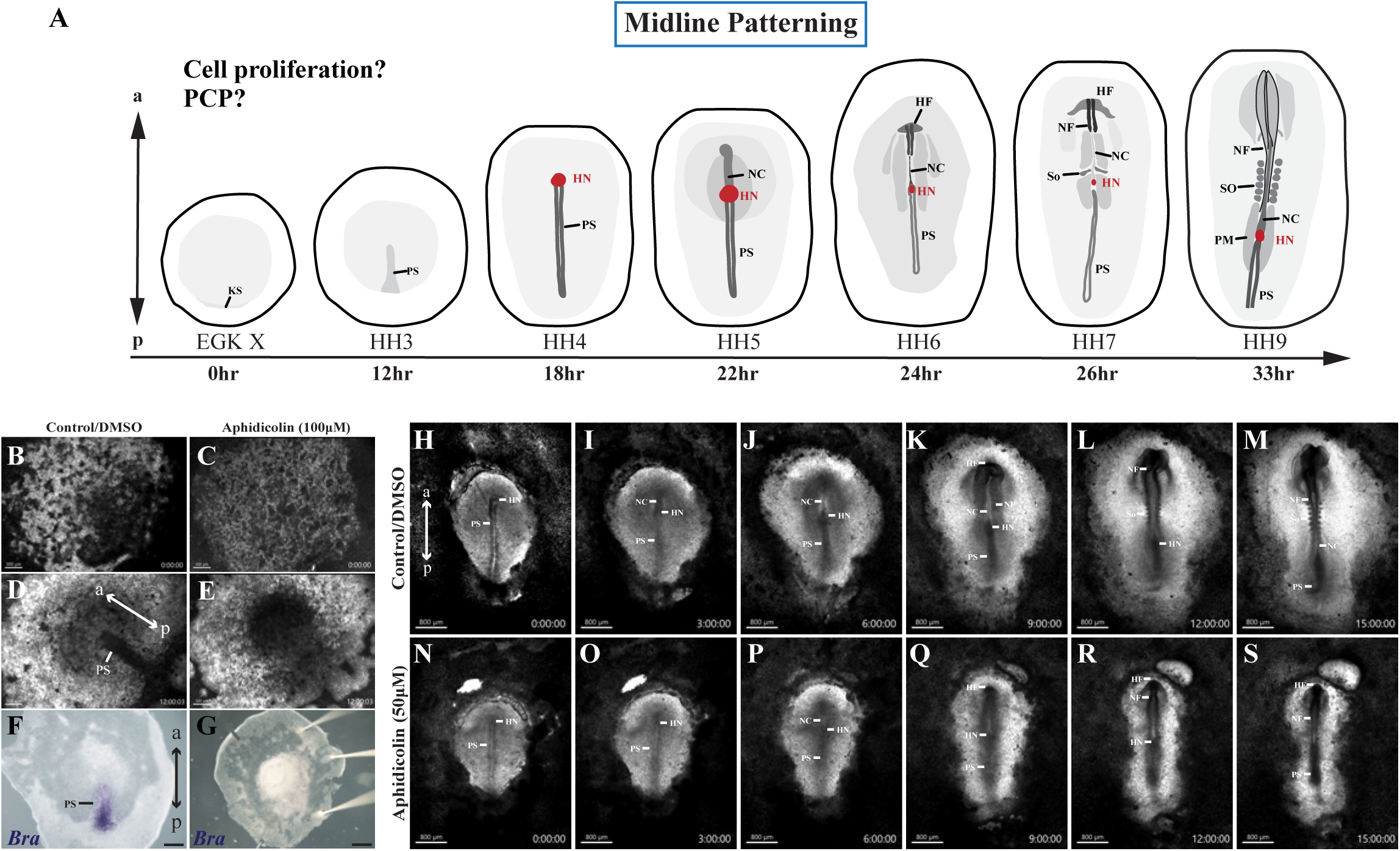
Formation of PS required cell proliferation, but midline patterning persisted upon aphidicolin treatment post PS-extension stage (HH4). Brightfield imaged embryos are viewed from the dorsal side (left side of the embryo corresponding to the reader’s left) with ap axis indicated by sign with two-end arrows. **(A)** schematic of chicken embryo development within the first 33hr after the egg has been laid with the question what role cell proliferation and PCP play in midline patterning. **(B, D)** Brightfield snapshots of DMSO-treated EGK X embryo that developed to HH3 with normal PS formation after 12hr. **(C, E)** Aphidicolin-treated EGK X embryo did not have PS formation upon 12hr incubation, scale bar = 300μm. **(F, G)** Presence or absence of PS marked by *Brachyury* expression in either DMSO- or aphidicolin-treated embryo, scale bar = 500μm. **(H-M)** Snapshots of every 3hr from timelapse of 15hr incubation after an HH4 embryo was treated with DMSO solution. HN appeared at the anterior tip of the fully extended PS, and then NC formed at the anterior of the HN when the HN/PS were regressing. Along with NC elongation and continuous regression of HN/PS, the HF formed at the anterior most of the embryo with visible neural plate which later turn into neural fold lateral to the axial structures. Somite formed after hr9, and embryo reached hh9 by hr15, Scale bar = 800μm. (N-S) Snapshots of every 3hr from timelapse of 15hr incubation after an HH4 embryo was treated with aphidicolin solution, scale bar = 800μm. KS. Koller’s sickle; PS, primitive streak; NC notochord; HN, Hensen’s node; HF, headfold; NF, neural fold; NP, neural plate; PM, paraxial mesoderm; So, somite.

Cell proliferation is present throughout embryogenesis and is vital for development and growth.^10,11,17–22^ The essential role of mitotic activities and PCP signaling in early amniote midline PS formation has been extensively studied in model animals, especially chick embryos given their easy accessibility for manipulations.^16,23,24^ Despite the importance, our understanding of the two distinct components, evolutionarily conserved and added, remains rudimentary for midline patterning after the full formation and extension of the PS, where HN forms at the anterior tip of the PS by Hamburger and Hamilton Stage 4 (HH4).^25,26^ After this stage, the post PS-extension phase of midline patterning is characterized by the HN shifting posteriorly together with the PS regression and shortening, leaving the anterior NC in its wake.^37,27^ Accompanied with HN regression and NC formation, the neural ectoderm (NE) and the neural fold (NF) subsequently form^27^ (Figure 1**A**, Movie S3).

Numerous mitotic Figures have been reported to occupy the chick neural plate, NC, and HN between stages HH7-12.^28,29^ Cell division within the NC proper as well as the addition of cells from the HN to the caudal side of the NC was proposed to be the main driving force for NC elongation,^29^ whether and how cell proliferation contributed to post PS-extension midline patterning was left to be further explored.

Convergent extension is a fundamental morphogenetic process underlies shape changes of various tissues in vertebrates and is regulated mainly through the PCP pathway.^9,30,31^ Core components of the PCP signaling network in vertebrates include the Frizzled (Fz) seven-pass transmembrane receptor, the four-pass transmembrane protein Van Gogh-like (Vangl), the Flamingo (Fmi/Celsr) cadherin with seven-pass transmembrane receptor features; the cytoplasmic protein Dishevelled (Dvl) recruited to the membrane by Fz, the cytoplasmic protein Prickle (Pk) recruited to membrane by Celsr, and the cytoplasmic protein Diego (Diversin/ankrd6/Inversin) recruited to membrane by Fz.^32^ They propagate PCP signal by asymmetric distribution within a cell where Fz-Dvl is on one side and Vang-Pk is on the other.^33^ Detailed observation and measurement of NC elongation demonstrates that the chick NC is getting narrower and longer – which fits the concept of convergent extension.^29^ However, whether and how PCP signaling might contribute to midline patterning after full PS-extension remains to be explored.

Here, we examined previously unexplored roles of both mitotic activities and PCP signaling for midline patterning both pre and post PS-extension in chick embryos. We report that midline patterning possesses two distinct phases to response to mitotic arrest: there was a severe loss of patterning and growth during PS extension, while in comparison patterning persisted when embryos were mitotically arrested post PS-extension. NC formation also persisted under mitotic arrest, but there was clear cell hypertrophy. Only a subset of surveyed PCP genes, including *prickle-1*, was detectable in midline structures at these post PS-extension stages, and morpholino-based Prickle-1 knockdown produced a modest effect on embryo and midline tissue lengths. In contrast to earlier periods with higher sensitivity to mitosis, later midline patterning relies less on cell proliferation and the PCP signaling pathway. These features put the midline patterning of amniotes in a unique place where they are only partially conserved with non-amniotes.

## Results

### Under mitotic arrest, PS extension is diminished, but midline patterning post PS-extension persists

At pre-streak and during PS-extension until HH3 stages, the cells of the epiblast and PS are highly proliferative,^11^ and mitotic arrest diminishes PS extension^12–14^ (Figure 1**B-E**, Movie S1- 2). During stages HH7-HH12, NC extension is driven in part by cell division within the NC proper as well as the addition of cells from the HN to the caudal side of the NC extension.^29^ However, the role of cell proliferation in midline patterning post PS-extension stages from HH4 up to stage HH7, which includes NC formation, HN/PS regression, and PS shortening, remains to be determined. To examine this, we first quantified mitotic activities by one-hour EdU incorporation at HH4 when the PS had fully extended along the midline. Our data indicated that cells in HH4 embryos were proliferative (Figure **S**1**A, B** and **D,** 50.00±6.98%, N=9), especially in the PS area (Figure **S**1**B’ and D**, 55.23±11.55%, N=8). In parallel, cells that were undergoing mitosis were detected by probing for phosphohistone 3, pH3^34,35^. Similar to EdU incorporation, there were higher portion of cells were in mitosis in the PS area (Figure **S**1**C’** and **E**, 5.14±0.65%, N=5). Both analyses of mitotic activities demonstrated that cells in HH4 chick embryos were proliferative.

To test whether these cell proliferation activities play any role in midline patterning from HH4 onward, whole embryos were exposed to Aphidicolin, which prevents cells from entering S-phase^36,37^; it has been widely utilized in the field to induce mitotic arrest in early chick embryos.^12–14^ Dynamic axial patterning events, such as NC formation as well as HN/PS regression, are slowed down by HH8-HH9.^38^ Embryos cultured *ex ovo* at these stages allow clear identification of the axial structures for various analyses. In our hands, control embryos cultured starting at stage HH4 reached stage HH8-9 after incubation times of 14-16hr on average, which led us to examine resulting embryos at 15hr hereafter. In contrast to the dramatic suppression of PS formation upon aphidicolin treatment (Figure 1**B**-**E**, Movie S2), embryos exposed to aphidicolin at HH4 continued to exhibit NC formation, HN/PS regression, as well as other morphogenetic and pattering events (Figure 1**N-S**, Table 1, and Movie S4) similar to the DMSO-only treatment control group (Figure 1**H-M**, Table 1, and Movie S3). Successful proliferation inhibition by aphidicolin was verified by a significant reduction in EdU incorporation (Figure **S**1**H, I,** and **J**, 0.002246±0.005265%, N=8, P<0.0001) compared to control (Figure S1**F, G** and **J**, 77.55±9.376%, N=5). There was also a reduction of cells in active mitosis upon aphidicolin treatment (Figure S1**H, I,** and **K**, 1.81±1.880%, N=5, P<0.1) compared to control (Figure S1**F, G** and **K**, 5.05±2.187%, N=3). No detectable change was observed in patterning the listed midline tissues upon initial examination. Our data suggest that axial patterning from HH4 can proceed under mitotic arrest until HH9.

**Table 1.** Summary of phenotypic analysis in embryos received DMSO/Aphidicolin treatment.

| Phenotype | No. of samples | DMSO |  |  | No. of samples | Aphidicolin |  |  |
| --- | --- | --- | --- | --- | --- | --- | --- | --- |
|  |  | Developed/Yes | Not develop/No | Abnormal |  | Developed/Yes | Not develop/No | Abnormal |
| Embryo development | 139 | 116 (83.45%) | 23 (16.55%) | 33 (23.24%) | 134 | 107 (79.85%) | 27 (20.15%) | 134 (100%) |
| Headfold formation | 139 | 88 (63.31%) | 51 (36.69%) | - | 134 | 59 (44.03%) | 75 (55.97%) | - |
| Neural fold/tube formation | 139 | 100 (71.94%) | 39 (28.06%) | - | 134 | 93 (69.40%) | 41 (30.60%) | - |
| NC formation | 139 | 130 (97.01%) | 9 (6.47%) | - | 134 | 107 (79.85%) | 27 (20.15%) | - |
| HN regression | 139 | 130 (97.01%) | 9 (6.47%) | - | 134 | 107 (79.85%) | 27 (20.15%) | - |
| Somite formation | 139 | 81 (58.27%) | 58 (41.73%) | - | 134 | 0 (0%) | 134 (100%) | - |
| PS cell aggregation | 139 | 1 (0.72%) | 138 (99.28%) | - | 134 | 123 (91.79%) | 11 (8.21%) | - |

### Midline identities are maintained upon mitotic arrest

The persistence of axial patterning programs upon mitotic arrest suggested that these tissues retain their corresponding identities. To address this possibility, we performed wholemount *in situ* hybridization (WISH) probing for various marker genes for midline tissues of interest. Markers used for NC identity included *Brachyury* (N=4), *Shh* (N=4), *Chordin* (N=6), and *Lefty-1* (N=3) which positively stained NC in DMSO/control embryos (Figure 2**A**, **C**, **E**, and **G**), while the tissue in the corresponding location was also labeled by the same transcripts in aphidicolin-treated embryos (Figure 2**B**, **D**, **F**, and **H**. *Brachyury*, N=4; *Shh*, N=5; *Chordin*, N=7; and *Lefty-1*, N=5). Markers for HN identity were *Chordin* (N=6) and *Nodal* (N=7) which detected the HN in both control embryos (Figure 2**E** and **Q**), and in aphidicolin-treated embryos (Figure 2**F** and **R**. *Chordin*, N=7; *Nodal*, N=8), indicating that regression of the HN took place in both conditions. PS identity was determined by probing for *Brachyury* (N=4) and *Fgf8* (N=7), both of which were detected in PS in control (Figure 2**A** and **I**) and aphidicolin-treated embryos (Figure 2**B** and **J**. *Brachyury*, N=4; *Fgf8*, N=8). Hybridization signals in aphidicolin-treated embryos were less intense and/or narrower; however, the detectable expression of marker genes in midline tissues in aphidicolin-treated embryos indicated that the midline continued to develop and pattern even when cell proliferation was inhibited.

**Figure 2.**
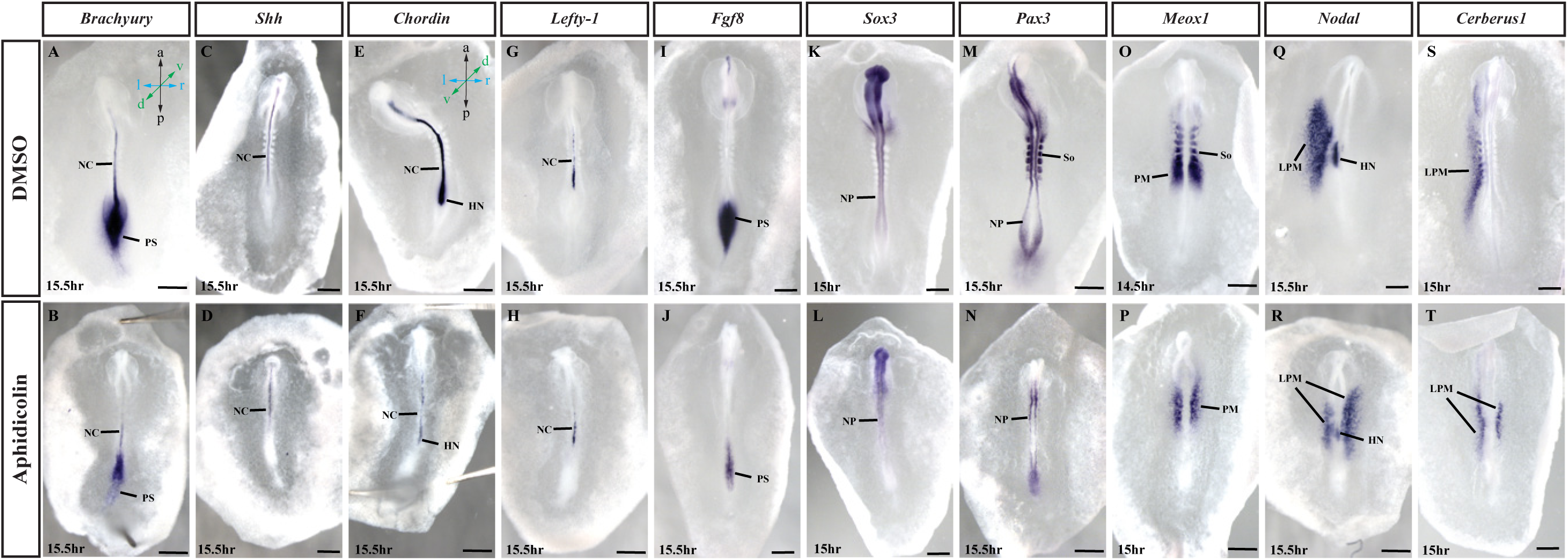
Gene expression analysis via WISH revealed midline identities and spatial expression pattern were maintained, but asymmetry was lost upon mitotic arrest. Wholemount in situ hybridization embryos in (**C-H)** are viewed from the ventral side (left side of the embryos corresponding to the reader’s right, indicated by the coordinate in (**E**). The rest of the embryos are viewed from the dorsal side (left side of the embryo corresponding to the reader’s left, indicated by the coordinate in (**A**). DMSO or Aphidicolin treatments are indicated on the left panel. Incubation time is indicated on the lower left of each image, in situ probe-targeted gene are indicated on the top panel, scale bar = 500μm. (**A**) Detection of *Brachyury* in DMSO and (**B**) Aphidicolin-treated embryos. (**C**) Detection of *Shh* in DMSO and (**D**) Aphidicolin treated embryos. (**E**) Detection of *Chordin* in control and (**F**) Aphidicolin treated embryos. (**G**) Detection of *Lefty-1* in DMSO and (**H**) Aphidicolin treated embryos. (**I**) Detection of *Fgf8* in control and (**J**) Aphidicolin treated embryos. (**K**) Detection of *Sox3* in DMSO and (**L**) Aphidicolin treated embryos. (**M**) Detection of *Pax3* in control and (**N**) Aphidicolin treated embryos. (**O**) Detection of *Meox1* in DMSO and (**P**) Aphidicolin treated embryos. (**Q**) Detection of *Nodal* in DMSO and (**R**) Aphidicolin treated embryos. (**S**) Detection of *Cerberus-1* in control and (**T**) Aphidicolin treated embryos. PS, primitive streak; NC, notochord; HN, Hensen’s node; NP, neural plate; PM, paraxial mesoderm; So, somite; LPM, lateral plate mesoderm.

To examine the patterning status of other embryonic tissues apart from the midline, we examined marker genes for the neural plate (NP), paraxial mesoderm (PM), somites (So), and lateral plate mesoderm (LPM). *Sox3* (N=8) and *Pax3* (N=6) transcripts were positive in the NP in both control (Figure 2**K** and **M**) and aphidicolin-treated embryos (Figure 2**L** and **N**, *Sox3*, N=9; *Pax3*, N=6), indicating the presence of NP under both conditions. *Pax3* was positive in somites in control embryos (Figure 2**M**), but it was negative in the corresponding somites area in the aphidicolin-treated embryos (Figure 2**M**). These results were consistent with brightfield images (Figure 1**L-S**, Movie S4) where somites were absent upon mitotic arrest, leading to the question of whether PM developed under such conditions. When we examined another somite marker, *Meox1,* which was expressed in somites (N=5) and PM (N=7) in control embryos (Figure 2**O**), it was also detected in aphidicolin-treated embryos at the area corresponding with either somite or PM (N=8) in the absence of segmented somites (Figure 2**P**). These data implied segmentation of the PM was diminished. Since Left-right asymmetry is established by this developmental window,^39^ we examined whether asymmetry establishment was preserved under mitotic arrest by probing for laterality marker genes. *Nodal* was expressed on the left side of the lateral plate mesoderm (N=12) and the HN (N=7) in control embryos (Figure 2**Q**); yet it was detected on both sides of the LPM (N=12) and the HN (N=8) in aphidicolin-treated embryos (Figure 2**R**), suggesting that asymmetry patterning was diminished under mitosis arrest. To verify this possibility, we examined another laterality marker gene, *Cerberus-1*. Of the total 11 embryos, 6 displayed a bilateral pattern in aphidicolin-treated embryos; 3 embryos displayed a bilateral pattern of larger expression domain on the left side compared to the right (Figure 2**T**) when compared to the left-sided expression in the LPM in control embryos (Figure 2**S**, N=8).

This result implied that cell proliferation might play a role in laterality preservation. The above findings suggest that axial tissues retained their identities when cell proliferation was inhibited – supporting the idea that midline patterning can proceed under mitotic inhibition post PS- extension stages. This led us to examine the underlying mechanism(s) that drives midline patterning under mitotic arrest, as seen in non-amniote embryos.

### Only a subset of PCP components was detectable in midline tissues

To address whether the midline structures especially NC formation and elongation under mitotic arrest are regulated by components of PCP signaling that are implicated in midline convergent extension in *Xenopus* embryos as well as PS extension in chicken embryos,^3,9,14,40^ we first performed WISH to survey gene expression of core PCP components. Although less quantitative, we chose this approach over other more quantitative methods, such as RT-PCR, for a better spatial resolution to specifically analyze the expression pattern of fine midline tissues.

Since WISH does not require surgical dissections of target tissues, thereby minimizes the potential complexity in interpreting outcomes due to contaminations of adjacent non-midline tissues.

Expression patterns of core PCP components in the axial structures were examined in post PS-extension stages HH4-8+ (5 somites) embryos, probing for *Van Gogh* (*Vangl2)*, *dishevelled* (*Dvl1* and *Dvl2/*3), *prickle (pk1)*, and *flamingo* (*Celsr1* and *Celsr2).* _3,14,40,41_ *Vangl2* was detectable in the PS and epiblast of a HH4 embryo (Figure 3**A**, N=5). It was detectable at headfold (HF), NP, PS, and the caudal side of the HN in HH5-HH7 embryos (Figure 3**B-D**, N=5, 4, 2). It was detected in NF and faintly at PS at HH8. However, by this developmental stage, it was no longer detectable at HN even after extended color development (Figure 3**E-F**, N=5). Consistent with a previous report,^42^ *Dvl1* was not detectable, or undistinguishable from the background, until HH8. Only a very faint staining signal, just above the background, was obtained in the PS even with extended color development (Figure 3**G-L**, N=4, 3, 5, 3, 5). *Dvl2/3* was detected in the epiblast, PS, HN, and mesoderm of an HH4 embryo (Figure 3**M**, N=4), and in NE, chordal-plate mesoderm, HN, and PS of stage HH5 embryos (Figure 3**N**, N=4). At stages HH6-7, *Dvl2/3* was still detected in HF, NP, PS, and the caudal side of HN (Figure 3**O-P**, N=5, 4). By HH8+ with 5 somites, a low signal became detectable in somites in addition to the above locations (Figure 3**Q-R**, N=4). A signal slightly above the background was detected in the NC in HH5-HH8 embryos even after prolonged color development. *Celsr1* was detected in the epiblast rostral of HN and the caudal most PS in HH4 embryos (Figure 3**Y**, N=8); and it was additionally detected in the NC faintly and NE in HH5-6 embryos (Figure 3**Z****-AA**, N=4, 3); on top of the above locations, it was enriched in NP/NF and LPM in HH7-8 embryos (Figure 3**AB****-AD**, N=3, 3). *Celsr2* was undetectable in HH4 embryos (Figure 3**AE**, N=5); and it was detected in NE/NP/NF in HH5-HH8 embryos (Figure 3**AF****-AJ**, N=5, 5, 4, 5). In addition, a weak signal was detected in the PS of HH8 embryos (Figure 3**AI-AJ**, N=5). *Pk1* was detected in the PS of HH4 embryos (Figure 3**S**, N=4). It was robustly enriched in NC-HN-PS area as well as PM in HH5-HH8 embryos (Figure 3**T-X**, N=4, 5, 6, 4), and faintly in somites starting from HH7 and in LPM of the same stages (Figure 3**X**, N=4). The major midline morphogenetic events from HH4-HH8 are NC formation/elongation, HN/PS regression, and PS shortening. However, WISH data of PCP core components indicated that only a subset of PCP signaling genes was expressed above the detectable level in the tissues of interest. The results raised the question as to whether the PCP signaling pathway was driving midline patterning during this developmental window.

**Figure 3.**
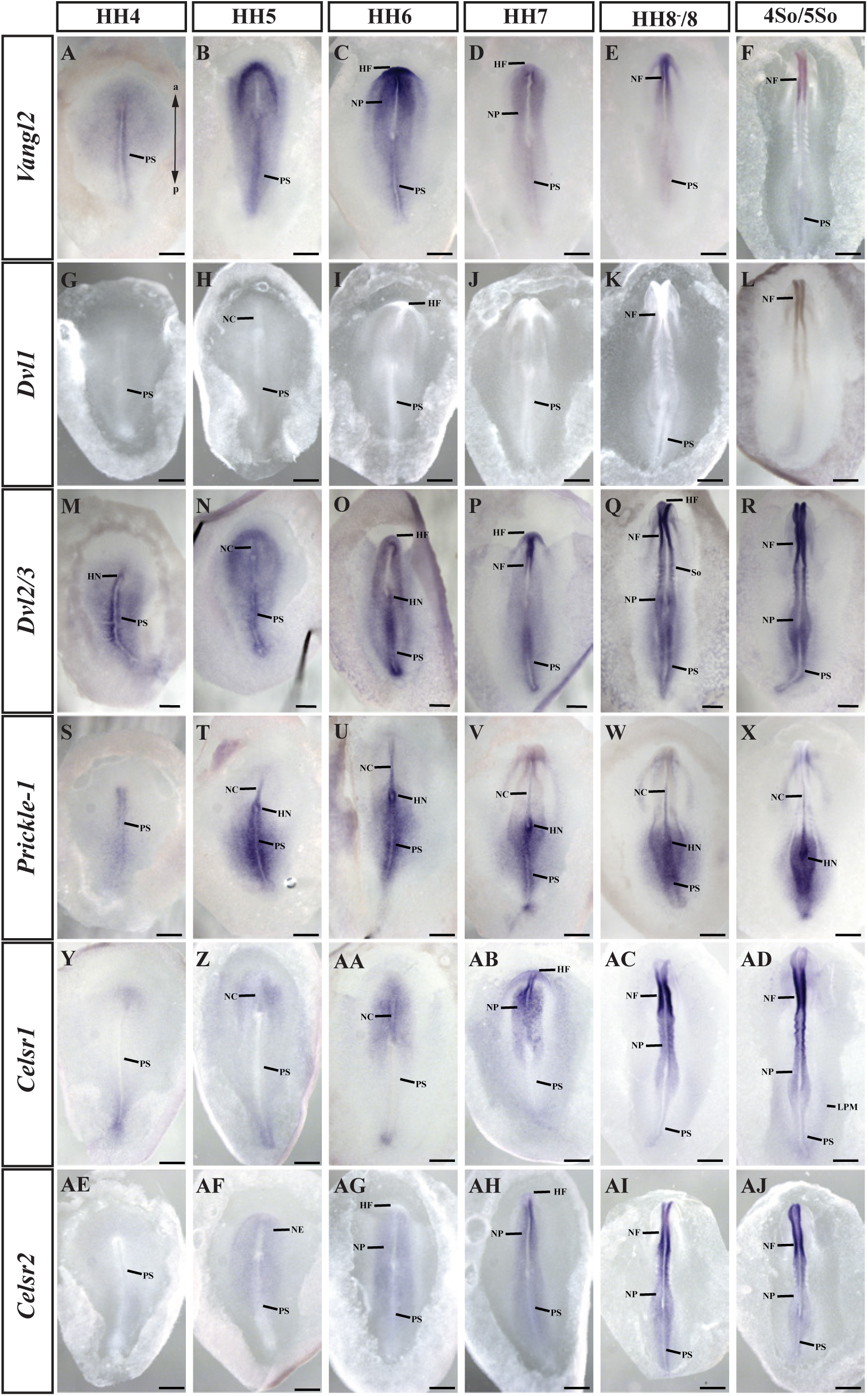
PCP core components expression analysis via WISH in HH4-HH8+ (5So) chick embryos. Embryos underwent WISH were viewed from the dorsal side (left side of the embryo corresponding to the reader’s left, anteroposterior axis indicated in (**A**). Targeted genes were indicated on the left panel while embryonic stages were indicated at the top panel, scale bar = 500μm. (**A**-**F**) Detection of *Vangl2* in HH4-5So embryos, N=5, 5, 4, 2, 5. (**G**-**L**) Detection of *Dvl1* in HH4-5So embryos, N=4, 3, 5, 3, 5. (**M**-**R**) Detection of *Dvl2/3* in HH4-5So embryos, N=4, 4, 5, 4, 4. (**S**-**X**) Detection of *Prickle-1* in HH4-HH8 embryos, N=4, 4, 5, 4, 4. (**Y**-**AD**) Detection of *Celsr1* in HH4-8 embryos, N=8, 4, 3, 3, 3. (**AE**-**AJ**) Detection of *Celsr2* in HH4-HH8 embryos, N=5, 4, 5, 6, 4. PS, primitive streak; NC, notochord; HN, Hensen’s node; HF, head fold; NE, neural ectoderm; NP, neural plate; NF, neural fold; So, somite; LPM, lateral plate mesoderm.

### Midline patterning persisted under Prickle-1 knockdown

To test the role of PCP signaling during midline patterning, we targeted *pk1* which was the only PCP component robustly detected in the NC-HN-PS area in our WISH-based survey (Figure 4**S-X**), which was aligned with the previous report.^43^ Morpholino-based knockdown was started at HH3+, before full PS-extension, and was incubated for 15hr. Fluorescently tagged control (cntl-MO) or Prickle-1 targeted (Pk1-MO) morpholino was introduced ubiquitously with midline-targeting into HH3+ embryos (Figure 4**B, C, F**, and **G**).

**Figure 4.**
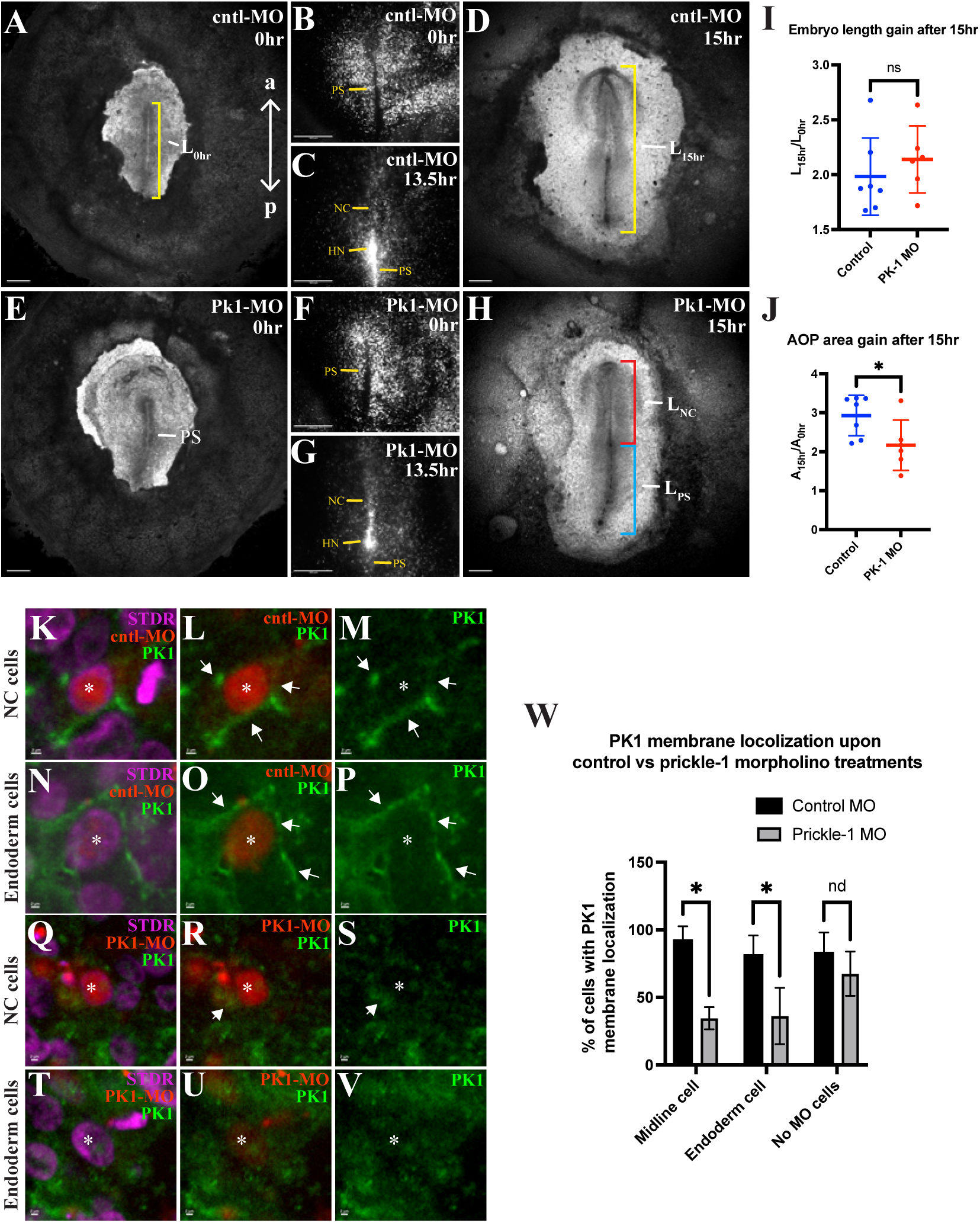
Midline patterning and embryo elongation did not deviate from control upon Prickle-1 knockdown, but embryonic area gain was reduced. (**A**, **E**) Brightfield images of HH3+ embryo immediately after introduction of cntl-MO or Pk1-MO; yellow bracket indicates embryo length at 0hr is represented by the length of PS, anteroposterior axis indicated by double-headed arrows. Scale bar = 500μm (**D**, **H**) Brightfield images taken upon 15hr of incubation; yellow bracket indicates length of embryo, red bracket indicates NC length, and blue bracket indicates PS length. Scale bar = 500μm. (**B**, **F**) Images showcasing ubiquitous and successful introduction of fluorescently tagged cntl- or Pk1-MO at 0hr and (**C**, **G**) at 13.5hr upon incubation, scale bar = 500μm. (**I**) Graph represents embryo length gain 15hr after MO introduction in both control and pk1-MO conditions, P=ns. (**J**) Graph represents embryo AOP area gain 15hr after MO introduction in both control and pk1-MO conditions, P<0.05. (**K**-**V**) Immunofluorescent images of NC cells and endodermal cells in either cntl-MO or Pk1-MO conditions. PK1, green; nuclear counterstain Sytox Deep Red, Magenta; morpholino, red. Asterixis indicate cells, white arrows indicate PK1. Scale bar = 2μm. (**W**) Graph represents PK1 membrane localization in midline cells, endoderm cells, and no MO cells in control and Pk1-MO conditions. Respectively, P<0.05, P<0.05, and P=0.1589, nd.

However, the resulting embryo with Pk1-MO displayed no readily detectable alterations in midline patterning or any other developmental defects distinguishable from the control (Figure 4**D** and **H**, cntl-MO, N=8; Pk1-MO, N=17). To quantify potential characteristics of the Pk1-MO induced perturbation, we examined the embryo length, as PCP signaling contributes to embryo elongation in many vertebrates.^31,44^ Since *Pk1* was mainly detected in the midline, we measured the length of the embryo at 0hr represented by the length of the PS and at the end of incubation represented by the length from HF to the caudal end of the PS.(Figure 4**A** and **D**, see **Experimental Procedures**) Our data showed that embryos reached similar lengths in both Pk1- MO-treated and control conditions (Figure S2**E**, cntl-MO, 3661±481.4μm, N=8; Pk1-MO, 3404±480.2μm, N=9, P=0.2882, ns). Moreover, embryo length gains for Pk1-MO-treated embryos were insignificant compared to control (Figure 4**I**, cntl-MO, 1.983±0.3519, N=7; Pk1- MO, 2.139±0.3049, N=6, P=0.4145, ns). Length gain was calculated by dividing embryo lengths at the end of incubation by lengths at 0hr (see **Experimental Procedures**). To test whether Pk-1 influences midline patterning, we measured the lengths of NC and PS at the end of incubation (Figure 4**H**). Neither NC nor PS lengths in Pk1-MO-treated embryos showed significant difference compared to control (Figure S2**F** and **G**, NC, P=0.1645; PS, P=0.4640, ns). Prickle-1 usually localizs to the cell membrane,^33,45^ and alteration of its localization could indicate successful knockdown. There was reduction of Prickle-1 membrane localization in both midline and the endoderm cells in embryos received Pk1-MO (Figure 4**Q**-**V**, and **W**, 34.63±8.23%, n= 154; 36.27±20.85%, n= 148, N=7, P<0.05) compared to control (Figure 4**K**-**P**, and **W**, 92.24±10.50%, n=92; 81.17±14.63% n=101, N=4). Membrane localization of Prickle-1 in those cells with no MO was comparable in both conditions (Figure 4**W**, cntl-MO, 82.97±15.15%, n=80, N=4; Pk1-MO, 67.68±16.38%, n=139, N=7, P=0.1589, nd). Hence, our data demostrated that Prickle-1 knockdown produced quantitative changes on its subcellular localization with only mild effect on embryo and midline tissue lengths. To test whether Pk-1 plays a role in overall embryo growth, we measured the area of the area pellucida (AOP) at 0hr and 15hr (Figure S2**A-D**). AOP areas of embryos that received Pk1-MO were smaller than control (Figure S2**H**, cntl- MO, 1.302*10^7^±2.823*10^6^μm^2^, N=8; Pk1-MO, 1.011*10^7^±1.598*10^6^μm^2^, N=9, P<0.05); moreover, Pk1-MO-treated embryos showed much smaller AOP area gain (Figure 4**J**, 2.168±0.6454, N=6) compared to control (2.926±0.5194, N=7, P<0.05). These data indicated that Pk1-MO introduction reduced overall embryonic growth. However, our data failed to demonstrate a potential contribution of Prickle-1 in midline patterning and embryo elongation. The role of Prickle-1 as well as PCP signaling during this developmental window remains to be explored.

### Persistent patterning along the midline upon mitotic arrest involved reducing tissue size and increasing cell size

Our morphological inspection and WISH data both demonstrated that multiple midline tissues formed with the expected corresponding identities despite inhibiting proliferation (Figure 1**N-S**, 2**B-N**, and **R**), indicating that the embryo continued differentiating and patterning along the midline. To summarize those morphogenetic events that persisted and those that diminished upon mitotic arrest, we undertook a phenotypic analysis based on anatomical features of tissues combined with molecular identities, using WISH analysis for tissue-specific marker genes (Figure 2). Detailed phenotypic analysis post PS-extension in anteroposterior order included patterning of HF, NF, NC, somite, HN, and PS (Table 1). Under our experimental conditions, both control embryos (83.45%, N=139) and those aphidicolin-treated (79.85%, N=134) survived and displayed midline patterning at a similar rate (Table 1). In contrast to persistent midline patterning, it was readily recognized that somites were absent under mitotic inhibition, which was aligned with our WISH data (Figure 2**N** and **P**, Table 1).

To gain insights into how embryos developed and patterned these midline tissues, we quantified size changes of the embryo and tissues in response to Aphidicolin-induced mitotic arrest. Aphidicolin-treated embryos were smaller based on brightfield image inspections (Figure 1**S**) and had narrower and weaker expression domains as seen through WISH (Figure 2**B-T**). To quantify the size change of the resulting embryos, both the area of AOP and the length of the embryo were measured at time zero and 15hr of incubation as described in previous section (Figure 5**A-D**, and **G-J, Experimental Procedures**). While control embryos demonstrated an area increase of over two-fold (Figure 5**B** and **E**, 2.345±0.512, N=124), aphidicolin-treated embryos grew less (Figure 5**D** and **E**, 1.289±0.337, N=112, P<0.0001). Control embryos displayed a nearly two-fold increase in length (Figure 3**F** and **H** 1.971±0.3064, N=105), and aphidicolin-treated embryos exhibited a significantly smaller increase (Figure 3**E** and **J**, 1.503±0.1939, N=97, P<0.0001). These results indicated that embryos at post PS-extension stages responded to proliferation inhibition by reducing embryo sizes while maintaining patterning along the midline.

**Figure 5.**
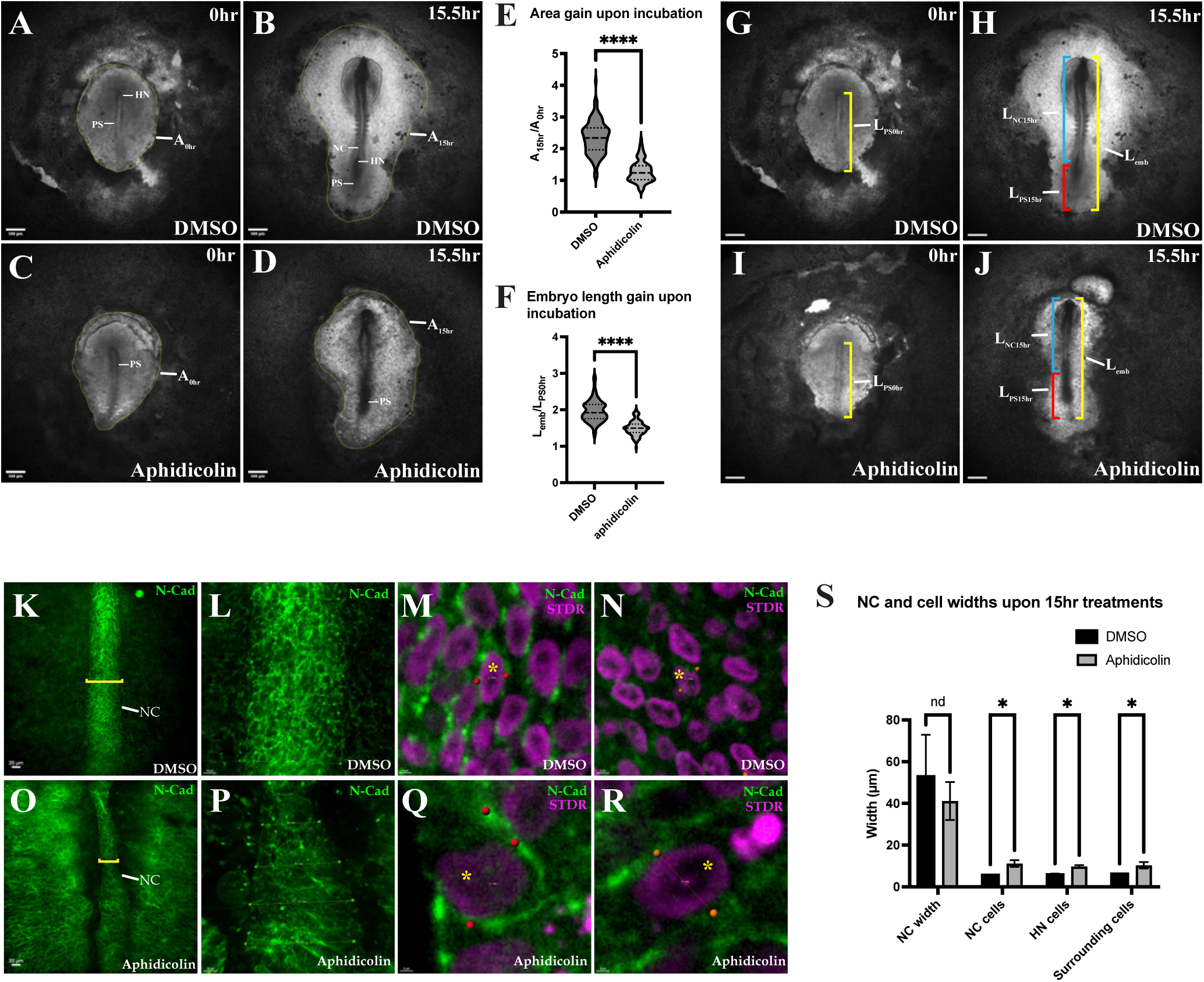
Chick embryos responded to proliferation inhibition with reduced area and embryo length gains together with narrower NC and widespread cell hypertrophy. (**A**, **C**) Brightfield snapshots from timelapse in which an embryo was either treated with DMSO/aphidicolin at 0hr and (**B**, **D**) at 15hr, yellow outlines indicate area pellucida (AOP), area measured as A_0hr_ and A_15hr_ respectively, scale bar = 500μm. (**E**) Graph compares area gain between DMSO and aphidicolin treatments; P<0.0001. (**G**, **I**) Brightfield snapshots from timelapse at 0hr and (**H**, **J**)15hr where an embryo was treated with DMSO/aphidicolin; yellow bracket indicates the embryo length at 0hr represented by PS length, L_PS0hr_, the embryo length at 15hr is L_emb_; blue bracket indicates the NC length at 15hr, L_NC15hr_; red bracket indicates PS length at 15hr, L_PS15hr_, scale bar, 500μm. (**F**) Graph compares embryo length gain between DMSO and aphidicolin conditions; P<0.0001. (**K**, **O**) Immunofluorescent staining images of NC area from control- and aphidicolin-treated embryos after 15hr incubation; N-cadherin, green. Yellow bracket indicates NC, scale bar = 20μm. (**L**, **P**) NC width measurements in Imaris, green spheres locate at the NC edges, and the distances between each pair of spheres represent the NC widths, P=ns, scale bar = 10μm. (**M**-**R**) Indicated by yellow asterisks, NC/surrounding cell widths measured in Imaris; red/orange spheres locate at the edges of the cell, and the distances between each pair of spheres measure the cell widths. Sytox Deep Red (STDR) labels nuclei, magenta, scale bar = 3μm. (**S**) Graph summaries NC and cell widths measured in control- and aphidicolin-treated embryos; P<0.0001 for all three groups. PS, primitive streak; NC notochord; HN, Hensen’s node.

Our morphological inspections of midline tissues revealed that aphidicolin-treated embryos continued with NC formation and HN/PS regression (Figure 1**N**-**S**, 5**D** and **J**, Table 1, Movie S4). Our length measurements revealed that aphidicolin-treated embryos exhibited a shorter length for both NC (1803±291.3μm, N=97, Figure S3**B**, P<0.0001) and PS (1299±269.8μm, Figure S3**C,** P<0.0001) with overall reduced AP-axis elongation (3120±334.8μm, Figure S3**A**, P<0.0001) compared to control (Figure S3**A**-**C**, NC, 2247±597.5μm; PS, 1670±481.0μm; Embryo, 3961±498.9μm; N=105). The compilation of measurements revealed that, under mitotic arrest, while the overall length of the embryo was decreased by 21.23%, the NC length was decreased by only 19.75%. Given our large sample sizes, the difference between these numbers prompted the question as to how, under mitotic arrest, the NC is preferentially lengthened within the embryo.

To address a potential mechanism, we examined the NC closely and measured its width via immunofluorescent staining against N-cadherin, which outlines cells in mesoderm and endoderm.^46^ Although cells were less organized in aphidicolin-treated embryos (Figure 5**O**), the NC was readily identified. The NC widths in aphidicolin-treated embryos were slightly reduced (Figure 5**P** and **S**, 41.14±9.09μm, N=8) when compared to controls (Figure 5**L** and **S**, 52.99±19.89μm, N=4, P=0.326, ns). After noticing enlarged cells in aphidicolin-treated embryos, we performed a cell size analysis. Our data revealed that cells of the NC became significantly larger in aphidicolin-treated embryos (Figure 5**Q** and **S**, 11.17±1.59μm, n=237, N=8) compared to control (Figure 5**M** and **S**, 5.69±0.298μm, n=391, N=4, P<0.0001). Enlarged cells were also found in the HN (Figure 5**S**, 9.75±0.72μm, n=78, N=7) as well as the area outside the NC in aphidicolin-treated embryos (Figure 5**R** and **S**, 10.35±1.52μm, n=448, N=8) while cells in the corresponding region in controls were significantly smaller (Figure 5**N** and **S**, 5.91±0.47μm, n=78; 6.23±0.09μm, n=352, N=4, P<0.0001). Although cell sizes were nearly doubled upon mitotic arrest, the ratios of NC and PS to the length of the embryo were similar in both aphidicolin-treated and the control conditions (Figure S3**D** and **E, ns**). Upon mitotic arrest, the chick embryo responded with embryonic size reduction together with cell hypertrophy; the latter may be a potential mechanism that contributed to persistent midline growth, but more importantly, preserved the midline proportionality and overall patterning.

## Discussion

Here, we have examined the role of cell proliferation in midline patterning of early amniote embryogenesis pre and post PS-formation. Contrasting to the highly mitosis sensitive nature of the PS formation^12–14^ (Figure 1**E**-**G**, Movie S2), post PS-extension embryos, when under mitotic inhibition, persisted with several midline patterning programs especially NC formation (Figure 1**N**-**S**, Table 1, Movie S4), which demonstrated that midline patterning in post PS-extension stages is less sensitive to mitosis. Hence, the data are consistent with the model wherein there are two distinct windows in midline patterning, with the transition from a reliance on mitosis lying in between stages HH3-HH4. While a non-amniote embryo pushed through development upon mitotic inhibition at late blastula, mid-gastrula, and early-neurula stages,^8^ we have shown that amniotes retain a facet of this for only a distinct period of development, after the PS has extended – similar to early-neurula. This contrasts with an amniote-specific requirement for mitosis during pre PS-extension stages.

We also investigated a role for PCP in midline patterning post PS-extension; These stages had not yet been examined. We presented data indicating that PCP might play a role in overall embryonic growth. Despite the insignificant alteration in embryo length gained as well as midline tissue lengths upon Prickle-1 knockdown (Figure 4**I**, S2**F,** and **G**), we cannot completely rule out PCP in midline patterning as well as embryo elongation. Therefore, the involvement of PCP in midline patterning remains unclear.

Although we surveyed the expression of several core PCP components expression in this current study, there are other homologues, as well as other components, were left to be surveyed.^32,33,47,48^ Due to the low-resolution nature of WISH, we cannot eliminate low transcript level of any components nor their expression status in the midline. *Prickle-1* was selected for its robust detection in the midline. *Dvl2/3* and *Celsr1* that were detected at low levels here (Figure 3**N** and **AA**) as previously reported,^41,42^ but they have yet to be tested for their role in midline patterning. Similarly, we cannot rule out functional redundancy of other *prickle* gene products. Prior to full PS-extension, a few PCP components have been examined at the protein level and have displayed polarization in PS as well as epiblast cells.^14,49^ However, PCP protein localizations within the midline cells post PS-extension were not examined except for Prickle-1. Our data have raised the question as to whether a positive role of PCP in post PS-extension midline patterning. A solution to this issue requires additional studies in the future.

Embryonic midline patterning includes but is not limited to cell proliferation and cell rearrangement. Our study focused on cell proliferation as well as the PCP signaling pathway, as for potential convergent extension of the NC. Thus, other important contributors, such as cell shape changes, intercellular space, extracellular matrix, and mechanical forces, were beyond the scope of our study and are left to be explored for their role in midline patterning. Embryonic size, as well as tissue length analysis upon mitotic arrest, revealed that embryos responded to a reduced cell population with reduced overall size while keeping midline tissue lengths proportional. Cell hypertrophy under mitotic arrest. alongside persistent NC elongation, led to our speculation that amniotes might utilize an evolutionary conserved mechanism that was initially discovered in fly imaginary disc^50–52^ where coordination between cell number and size work together to maintain tissue/organ size. However, lacking the total cell number of the entire embryo as well as the NC and other tissues, limits our understanding of how cell proliferation and cell size contribute to midline patterning. Furthermore, cell generation time in chordamesoderm rostral to the HN in an HH5 embryo has not been measured, but rather calculated to be 7-10hrs.^10^ That means our effective cell proliferation inhibition can best be 1-2 cell cycles, which leads to the question of where the supply of cells is from, upon mitotic arrest. We cannot rule out incomplete inhibition, yet there might be alternative explanations. The HN is a well-known stem cell containing enlarged structure in stage HH4 embryos^25,53–55^; it is reasonable to theorize that there might be enough resident cell population to support both caudal regression of the HN and early phase elongation of the NC. Reduced WISH signal in both NC and HN upon mitotic arrest (Figure 2**B**, **D**, **F,** and **H**) could support such theory but requires additional evidence.

Our study has established an atlas showcasing the roles of cell proliferation and PCP in midline patterning pre and post PS-extension using chick embryos. The two mitotic sensitivity phases of midline patterning put the amniote in an evolutionary unique place distinguished from non-amniotes. Our study provides a fundamental understanding of the role of cell proliferation in midline patterning but also attracts attention to the rarely explored initial phase of NC formation and HN/PS regression. Our work would inspire multiple potential directions, in addition to cell proliferation and PCP pathway targets, for studying amniote midline patterning as well as other developmental programs. Some new questions presented, such as how cell proliferation contributes to laterality establishment/maintenance or segmentation of the PM, would be valuable additions to our current knowledge.

### Experimental Procedures

#### Embryo isolation and *ex ovo* culture

Fertilized eggs of White Leghorn (*Gallus Gallus domesticus*) were obtained from Petaluma Farms (Petaluma, CA) and Avis Bio (Norwich, CT), and incubated at 38.3°C in a humidified incubator to desired embryonic stages.^26^ Both dissection methods with and without filter paper ring are performed as previously published.^56–58^ Embryos were isolated in Tyrode’s solution (137mM NaCl, 2.7 mM KCL, 1mM MgCl2, 1.8mM CaCl2, 0.2 mM Ha2HPO4, 5.5mM D-glucose, pH 7.4). After manipulations (inhibitor application or electroporation), embryos were incubated or live-imaged ventral side up on a vitelline membrane stretched around a custom made glass ring according to the New culture method as previously described.^59^

### Aphidicolin/DMSO treatment on ex *ovo* culture

Pre-streak embryos were dissected and treated with DMSO/Aphidicolin as previously described.^14,58^ Post PS-extension embryos were dissected via filter paper method^57^ using a piece of concentric circular Whatman filter paper with 0.65in outer diameter and 0.15in inner diameter. Embryo culture media was a mixture of 5:1 of albumin and Tyrode’s solution that contained DMSO (0.17% v/v) or aphidicolin (50μM). After the embryos were laid down with ventral side up, 20μL of the same concentration of DMSO or aphidicolin in Tyrode’s solution were applied over the embryos. Embryos were then sent to imaged or incubated as described.

### EdU assay

10mM of EdU stock was diluted into Tyrode’s solution that contained DMSO/aphidicolin to reach a final concentration of 100μM; this working solution was applied directly over the embryos prior to imaging or incubation. For measuring proliferation rate in stage HH4 embryos, the EdU stock was instead diluted in 1X PBS (Gibco™, 14190250) and 10μL of 100μM final working solution was gently applied over the embryos and incubated for 1hr at 38°C. After incubation, embryos were fixed with 4%PFA (Electron Microscopy Science, 15710) in 1X PBS for 30min at RT, then permeabilized with 0.5% Triton™ X-100 (Sigma-Aldrich, T8787) in 1X PBS for 30min at RT. EdU signal was detected by following the manufacturer protocol for cells (Invitrogen, C10338). Immunofluorescence staining and nuclear counter stain were performed afterward as described in the following section.

### Immunofluorescence staining

Embryos were fixed with 4%PFA/PBS for 30min at RT, and then embryos were washed 3 times with 1X PBS before being permeabilized and blocked with 5% normal goat serum (Abcam Inc, ab7418) in 0.1% PBST (Triton™ X-100) for 2hr at RT. The solution was replaced with primary antibodies diluted in 5% GS/0.1% PBST to incubate with gentle rocking overnight at 4°C. Primary antibody used include anti-phosphohistone-3 antibody (mouse 1:1000, Cell Signaling, 9706, Clone 6G3), anti-N-cadherin antibody (mouse 1:200, Sigma-Aldrich, C3865, Clone GC4), and anti-Prickle-1 antibody (rabbit, 1:300, Proteintech, 22589-1-AP). After 3X washes with 0.1%PBST at RT, the solution was replaced by secondary antibodies that were diluted in 5% GS/0.1% PBST to incubate with gentle rocking overnight at 4°C (or 2-3hr at RT). Secondary antibodies used include: goat anti-mouse IgG (H+L) Alexa Fluor™ 488 (1:500, Invitrogen, A- 11029) and goat anti-rabbit IgG (H+L) Alexa Fluor™ 488 (1:500, Invitrogen, A-11008). Nuclei were labeled by SYTOX™ Deep Red Nucleic Acid Stain (1:1000, Invitrogen, S11380) together with secondary antibodies. After 2X 30min washes with 0.1% PBST at RT, the embryos were equalized and mounted with refractive index matching media (RIMs, 80% w/v Histodenz**^™^**, Sigma-Aldrich, D2158; 0.01% sodium azide, 0.1% Tween 20, pH7.5).(^60^ All samples mounted in RIMs were imaged within a week to retain the best clearing effect and imaging quality.

### Whole-mount *in situ* hybridization (WISH) and subsequent immunohistochemistry

Probes were generated from plasmids for *Brachyury* and *Sox3* (gifts from Dr. Raymond B. Runyan), *Fgf-8,*^61^ *Nodal, Lefty-*1, and *Shh*^62^; *Pax-3* (Clone OGa25640), *Meox-1* (Clone OGa25554), and *Cerburus-1* (Clone OGa26937) custom ordered from GenScript; *Chordin,*^16^ *Vangl2,*^15^ *Prickle-1,*^43^ *Celsr-1*, *Celsr-2,*^63^ *Dishevelled-1, and Dishevelled-2/3.*^42^ Embryos were fixed with 4%PFA/PBS for 30min at RT or overnight at 4°C. WISH protocol was adapted from both Geisha ^63^ and previous published protocol from our lab.^14,56^ All WISH were carried out with paralleled control embryos for managing the color development time. Embryos were washed 3X with 0.1%PBST (Tween^®^ 20, Sigma-Aldrich, P1379) for 10min at RT, and then dehydrated with graded methanol/PBS solution and ended with 100% methanol. After O/N in 100% methanol at - 20°C (Day 1), the embryos underwent graded rehydration and followed by washes with 1XPBS. The embryos were treated with 5μg/mL proteinase K/PBS (Invitrogen, AM2546) for 1min at RT and followed by several washes with 0.1%PBST. The embryos were then refixed with 0.2%glutamialdehyde/4%PFA/PBS for 30min at RT, and washed 3X 5min with 0.1%PBST. After the embryos were sunk in prehybridization solution [10mL Solution 1 (1% SDS, 5X SSC, 50% formamide), 10μL of heparin (50μg/mL), 50μL yeast RNA (50μg/mL)], they were incubated in freshly replaced prehybridization buffer for 3hr at 70°C. Meanwhile, the hybridization buffer that contained 10μg/mL targeted DIG-labeled RNA probes was warmed up for 3hr at 70°C. The prehybridization buffer was then replaced with hybridization buffer and the embryos were hybridized O/N at 70°C. (Day 2) Embryos were washed 3x20min with Solution 1 at 70°C, and then followed by 3x20min washes with freshly made Solution 3 (50% formamide and 2X SCC pH 4.5) at 62°C. Then, the embryos were washed with Tris-Buffered Saline (pH7.5) with 0.1% Tween 20 (TBST) 3x20min at RT and incubated with blocking solution (10% sheep serum in TBST) for 2hr at RT. Anti-DIG-AP Fab fragment antibody (1:2000, Roche, 11093274910) in blocking solution was applied to embryos and incubated O/N at 4°C with gentle rocking. (Day 3) Embryos were washed in TBST at least 5x1hr at RT and then O/N washed at 4°C with gentle rocking. (Day 4) Embryos were washed in freshly made NTMT (100mM NaCl, 100mM Tris, pH9.9, 50mM MgCl_2_, 0.1% Tween 20) for 15min at RT, and then replaced with color-developing reagent: 2.3μL NBT (Roche, 11383213001) and 3μL BCIP (Roche, 11383221001) diluted in 1mL NTMT and incubated in the dark at RT or 4°C until appropriate signal developed. Reactions were stopped by adding and washing with TBST, then embryos were washed several times with PBS followed by placing them in 4% PFA/PBS at 4°C for long term storage.

### Electroporation

Embryos were transfected with Prickle-1 morpholino, or control-oligo DNA conjugated with Lissamine (GENE TOOLS) using an electroporator (Nepagene) with 3 pulses of 5-5.5 Volts, 50 milliseconds duration, 500 milliseconds interval, and platinum electrodes. Approximately 1.37μL of morpholino solution was delivered to the embryo: Morpholino solution contained 1% fast green (final 0.27%), 80% glucose (final 14%), 5μg/μL of H2B-BFP expression vectors (final 1.36μg/μL), and 1mM Prickle-1 morpholino (5’- TTGTTAGCTTTGGGCTCCATCTCCA 3’- Lissamine) or control-oligo (5’-CCTCTTACCTCAGTTACAATTTATA-3’-Lissamine) (final 0.27mM).

### Imaging of chick embryo

For live imaging, embryos were cultured in a 35mm dish with New culture method at 38°C. Time-lapse images were recorded with Nikon Widefield Epifluorescence inverted microscope (Nikon Eclipse TE2000-E supported by Hamamatsu ORCA-Spark camera with Nikon Plan Apo 2x/0.1 objective). The acquisition time was every 3 minutes using Nikon Elements Advance Research software V4.00.07.

Embryos that were successfully electroporated with fluorescently tagged morpholino were imaged by Echo Revolve R4 inverted microscope with Olympus UPLANFL N 4x PHL objective.

Immunofluorescence-stained chick embryos were imaged by a Crest X-Light V2 L-FOV spinning disk confocal on Nikon Ti2-E Microscope with Prime 95B 25mm sCMOS camera at the Center for Advanced Light Microscopy at UCSF. EdU incorporated and pH3 stained embryos were scanned with CFI Plan Apo Lambda 20x/0.75 lens with 0.92μm z-step size and 20% overlapping area, and then images were stitched using Nikon Elements Advance Research software V5.2. NC-containing areas in N-Cadherin and Prickle-1 stained embryos were imaged with CFI Plan Apo Lambda S 40x/1.25 Corr (Silicone) lens with 0.33μm z-step size. Fixed WISH samples were imaged by Leica MZ16F microscope with Leica DFC300 Fx camera and Leica FireCam V.3.4.1 software.

### Quantification and statistical Analysis

#### Cell number, EdU-assay, pH3 detection, and mitotic rate

All the measurements within this section were done with Imaris Spot Detection tool.

For HH4 embryos, multiple cuboids that were 165μm*165μm*z were created to survey the PS, lateral areas on both the left and right side of the PS and the anterior area to the PS. The number of cells, EdU-positive cells, and pH3-positive cells were measured within these cuboids. The mitotic rate was calculated by dividing either the number of EdU- or pH3-positive cells by the total number of cells measured in an embryo as a single biological sample. Midline cells were represented by areas selected along the PS (Figure S1**A’**, **B’** and **C’**), the anterior cells were represented by selected areas anterior to the PS, lateral left/right cells refer to areas selected left/right lateral to the PS, overall refers to the sum of all the above (Figure S1**D-E**).

For aphidicolin/DMSO-treated embryos, the same cell number detection process was performed. Surveyed areas within embryos were kept consistent between aphidicolin and DMSO conditions to reflect the effectiveness of mitotic inhibition. The overall mitotic rate was calculated the same as HH4 embryos mentioned above (Figure S1**J** and **K**)

#### Embryo area and various length measurements

Embryo area measurement was represented by area pellucida (AOP), manually outlined in ImageJ by Freehand selections function at the timepoint of interest (Figure S2**A-D**, 5**A-D**). The embryo area gain was calculated via embryo area at the end of incubation divided by area measured at 0hr of incubation. At 0hr/HH4, embryo length was represented by the length of PS manually traced in ImageJ with Segmented Line function; at the end of incubation, embryo length was represented by the length from the anterior end of headfold to the posterior end of PS manually traced with Segmented Line function. Embryo length gained/changed was calculated by dividing embryo length at the end of incubation (usually 15hr) by the length of PS at 0hr. At the end of incubation, the length of NC is represented by the length from the anterior end of the headfold to HN manually traced by Segmented Line, and the length of PS is represented by the length from HN to the posterior end of the PS traced with Segmented Line. Embryo length equals the sum of NC and PS lengths. NC proportion was calculated by dividing NC length by embryo length. PS proportion was calculated via dividing PS length by embryo length.

#### Notochord and cell width measurements in aphidicolin and DMSO treatment embryos

Embryos that were treated with DMSO/aphidicolin underwent immunofluorescent staining of N-cadherin, which is widely expressed in mesoderm, for measuring notochord and cell widths. Both NC and cell width measurements were performed in Imaris software with Measurement Tool. Lines parallel to left-right axis were created by pairs of two points at the left and right edges of NC, indicated by N-cadherin staining, and were used to measure the width of the NC (Figure 5**L** and **P**). Multiple widths along the AP axis were measured within one embryo to get an average width for one biological sample. The width of cells at various locations was measured by the distance between two points across the narrowest part of a cell (Figure 5**M-R**).

#### Statistics

Statistical analyses were performed using two-tailed unpaired Student’s t-test and unpaired t-test with Welch’s correction in Prism. Individual P values and sample numbers were described in the text, Figure panels, and legends. Graphs were generated using GraphPad Prism 10.2.2 for dot plots, bar plots, histograms, and violin plots. Error bars in all figures represent the standard deviation of the mean (SD). P<0.05 and P<0.1 were considered statistically significant. *P < 0.1 or 0.0332, **P < 0.0021, ***P < 0.0002, ****P < 0.0001.

## Acknowledgements

We thank past and present Mikawa lab members, particularly Dr. Jeanette Hyer for her invaluable suggestions and/or assistance; we also thank Buckwalter lab for providing the Echo Revolve microscope for image acquisition. Imaging data for this study were acquired at the Center for Advanced Light Microscopy at UCSF on CREST/C2 Confocal obtained using grants from the UCSF Program for Breakthrough Biomedical Research funded in part by the Sandler Foundation and the UCSF Research Resource Fund Award. This work was supported in part by grants from NIH (R01HL153736, R01HL148125) to T.M.; Uehara Memorial Foundation Fellowship and JSPS Postdoctoral Fellowship for Research Abroad to R.A. Eunice Kennedy Shriver National Institute of Child Health and Human Development (NICHD) Predoctoral Training in Developmental Biology T32HD007470 to Z.Z.; it is part of Z.Z.’s PhD thesis work.

## Author contributions

T.M., Z.Z., and R.A. conceptualized the project. T.M., Z.Z., and R.A. wrote the manuscript. Z.Z. and R.A. designed and performed all experiments. Z.Z. and R.A. carried out all the statistical analysis.

