## supplemental figures for "Differential Sensitivity of Midline Patterning to Mitosis during and after Primitive Streak Extension"

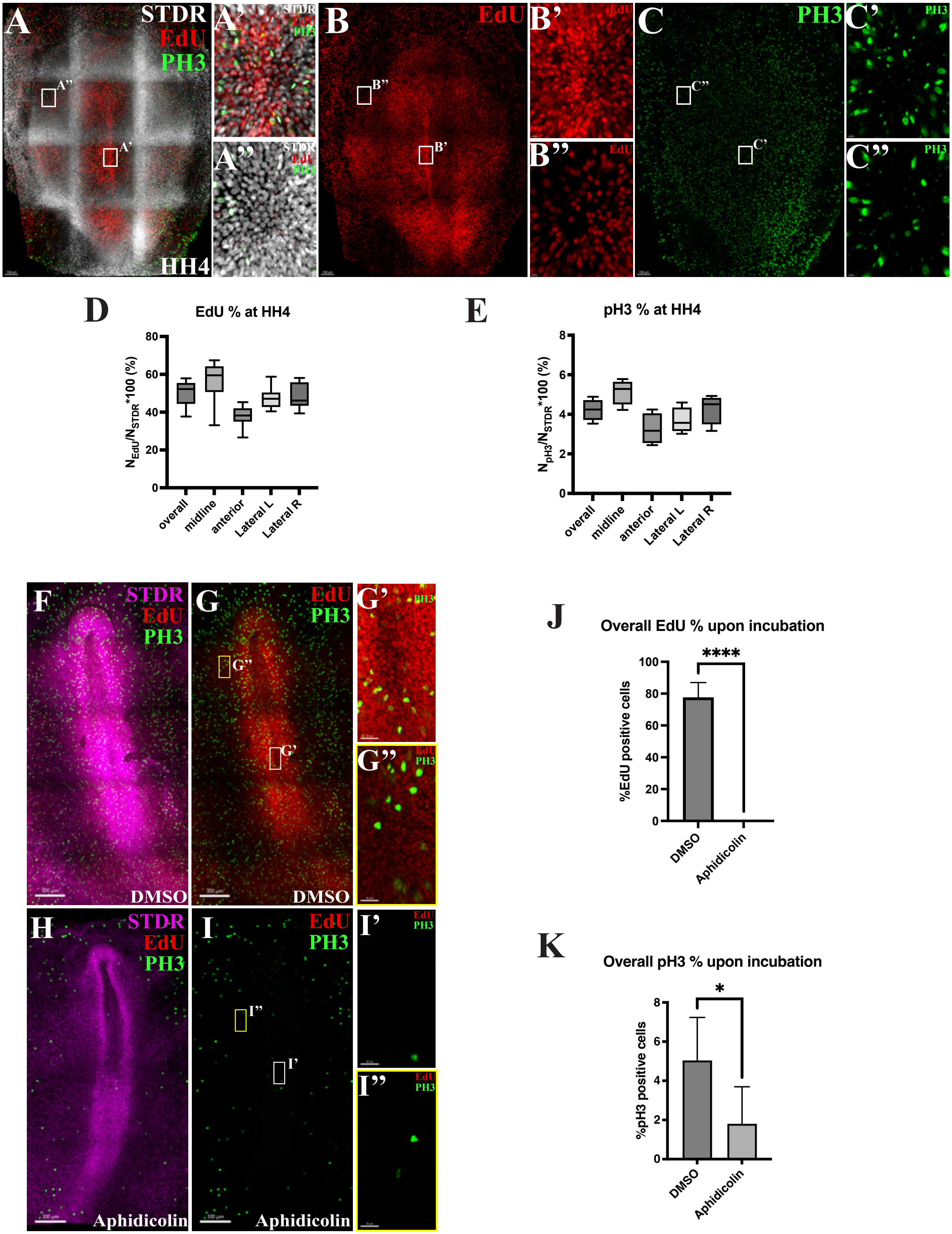

A
STDR
EdU
PH3
HH4
A'
A''
B
EdU
B'
B''
C
PH3
C'
C''
D
EdU % at HH4
$N_{EdU}/N_{STDR}$*100 (%)
overall
midline
anterior
Lateral L
Lateral R
E
pH3 % at HH4
$N_{pH3}/N_{STDR}$*100 (%)
F
G
G'
G''
DMSO
H
I
I'
I''
Aphidicolin
J
Overall EdU % upon incubation
%EdU positive cells
****
DMSO
Aphidicolin
K
Overall pH3 % upon incubation
%pH3 positive cells
*

**Figure S1. Cells were proliferative in HH4 embryos, and inhibition of cell division by aphidicolin was effective.**

(**A-B''**) EdU (red) incorporation and (**C-C''**) pH3 (green) staining in HH4 embryos; nuclear counterstain by Sytox Deep Red (STDR, magenta), scale bar = 150µm. (**A', B', and C'**) Enlarged view of the midline and (**A'', B'', and C''**) lateral left, scale bar=10µm. (**D**) Percentage of EdU positive cells in HH4 embryos; overall, 50.00±6.984%, n=114138; midline, 55.23±11.55%, n=39566; anterior, 37.85±6.214%, n=15439; lateral left, 47.55±6.285%, n=30280; lateral right, 48.90±7.26%, n=28853, N=7. (**E**) Percentage of pH3 positive cells in HH4 embryos; overall, 4.225±0.5553%, n=68519; midline, 5.142±0.6548%, n=22024; anterior, 3.258±0.7948%, n=11012; lateral left, 3.691±0.6613%, n=185532; lateral right, 4.279±0.7717%, n=16951, N=4. (**F-G**) EdU incorporation (Red) in DMSO and (**H-I**) aphidicolin treated embryos, which are stained with pH3 (green) and STDR (magenta), scale bar = 300µm. (**G' and I'**) Enlarged view of the midline and (**G'' and I''**) lateral left indicated in, scale bar = 30µm. (**J**) Overall percentage of EdU positive cells in DMSO- and aphidicolin-treated embryos, DMSO, 77.55±9.376%, n=671730, N=5; aphidicolin, 0.002246±0.005265%, n=683816, N=8, P<0.0001. (**K**) Overall percentage of pH3 positive cells in DMSO- and aphidicolin-treated embryos, DMSO, 5.052±2.187%, n=83346, N=3; aphidicolin, 1.815±1.880%, n=59028, N=5, P<0.1.

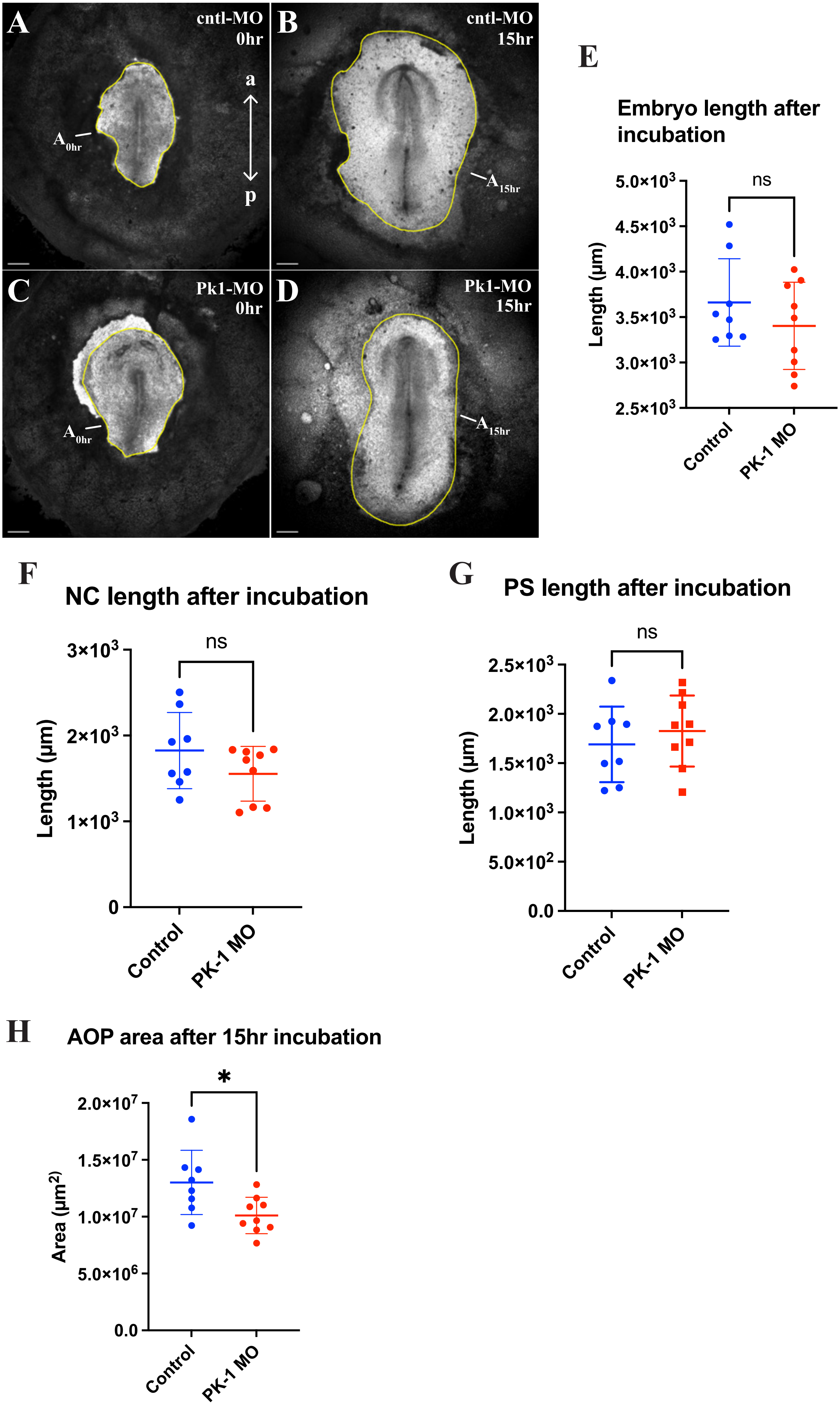
A
cntl-MO
0hr
a
p
$A_{0hr}$
B
cntl-MO
15hr
$A_{15hr}$
C
Pk1-MO
0hr
$A_{0hr}$
D
Pk1-MO
15hr
$A_{15hr}$
E
Embryo length after incubation
ns
Length (µm)
5.0×10^3
4.5×10^3
4.0×10^3
3.5×10^3
3.0×10^3
2.5×10^3
Control
PK-1 MO
F
NC length after incubation
ns
Length (µm)
3×10^3
2×10^3
1×10^3
0
Control
PK-1 MO
G
PS length after incubation
ns
Length (µm)
2.5×10^3
2.0×10^3
1.5×10^3
1.0×10^3
5.0×10^2
0.0
Control
PK-1 MO
H
AOP area after 15hr incubation
*
Area (µm^2)
2.0×10^7
1.5×10^7
1.0×10^7
5.0×10^6
0.0
Control
PK-1 MO

**Figure S2. Midline patterning upon Pk1 knockdown remained comparable to control, but embryo area was reduced.**

(**A**-**D**) Brightfield images of embryos treated with control- or PK1-morpholino at 0hr and 15hr, yellow outline indicates area pellucida (AOP), scale bar = 500μm. (**E**) Graph compares embryo length at 15hr post treatment of cntl- and PK1-MO, related to Figure 4**D** and **H**; cntl-MO, 3661±481.4μm, N=8; PK1-MO, 3404±480.2μm, N=9; P=0.2882, ns. (**F**) Graph compares NC length at 15hr post treatment of cntl- and PK1-MO, related to Figure 4**D** and **H**; cntl-MO, 1826±443.4μm, N=8; PK1-MO, 1554±318.6μm, N=9; P=0.1645, ns. (**G**) Graph compares PS length at 15hr post treatment of cntl- and PK1-MO, related to Figure 4**D** and **H**; cntl-MO, 1690±383.7μm, N=8; PK1-MO, 1826±360.4μm, N=9; P=0.4640, ns. (**H**) Graph compares AOP area measurements, cntl-MO, $1.302*10^7 \pm 2.823*10^6 \mu m^2$, N=8; PK1-MO, $1.011*10^7 \pm 1.598*10^6 \mu m^2$, N=9; $P<0.05$.

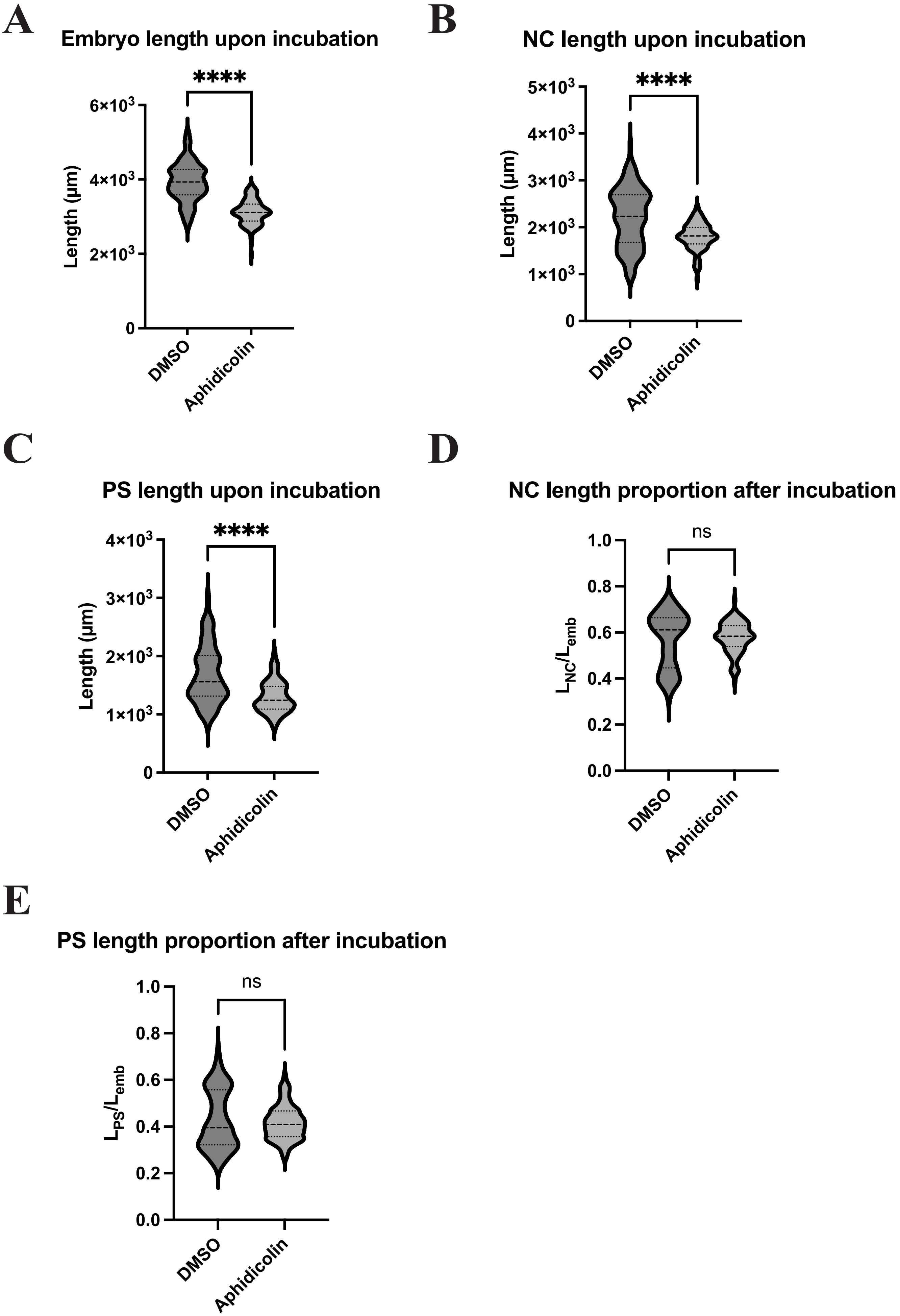
A
Embryo length upon incubation
****
6×10^3
4×10^3
2×10^3
0
Length (µm)
DMSO
Aphidicolin
B
NC length upon incubation
****
5×10^3
4×10^3
3×10^3
2×10^3
1×10^3
0
Length (µm)
DMSO
Aphidicolin
C
PS length upon incubation
****
4×10^3
3×10^3
2×10^3
1×10^3
0
Length (µm)
DMSO
Aphidicolin
D
NC length proportion after incubation
ns
1.0
0.8
0.6
0.4
0.2
0.0
L_NC/L_emb
DMSO
Aphidicolin
E
PS length proportion after incubation
ns
1.0
0.8
0.6
0.4
0.2
0.0
L_PS/L_emb
DMSO
Aphidicolin

**Figure S3. NC, PS and embryo length were significantly shorter upon mitotic arrest compared to control, but NC and PS length proportions remained similar between the two conditions.**

All measurements were taken at 15hr post incubation in DMSO- or aphidicolin-treated embryos, related to Figure 5**H** and **J**. All graphs compare measurements between DMSO and aphidicolin conditions. (**A**) Embryo length, DMSO, $L_{emb}$ = 3961±498.9μm, N=105; aphidicolin, $L_{emb}$ = 3120±334.8μm, N=97, P<0.0001. (**B**) NC length, DMSO, $L_{NC}$ = 2247±597.5μm, N=105; aphidicolin, $L_{NC}$ = 1803±291.3μm, N=97, P<0.0001. (**C**) PS length, DMSO, $L_{PS}$ = 1670±481.0μm, N=105; aphidicolin, $L_{PS}$ = 1299±269.8μm, N=97, P<0.0001. (**D**) NC length proportion to embryo, DMSO, $L_{NC}/L_{emb}$ = 0.5673±0.1203, N=105; aphidicolin, $L_{NC}/L_{emb}$ = 0.5788±0.06670, N=97, P=0.3995, ns. (**E**) PS length proportion to embryo, DMSO, $L_{PS}/L_{emb}$ = 0.4230±0.1213, N=105; aphidicolin, $L_{PS}/L_{emb}$ = 0.4162±0.07330, N=97, P=0.6296, ns.
