## supplemental movie 1 for "Differential Sensitivity of Midline Patterning to Mitosis during and after Primitive Streak Extension"

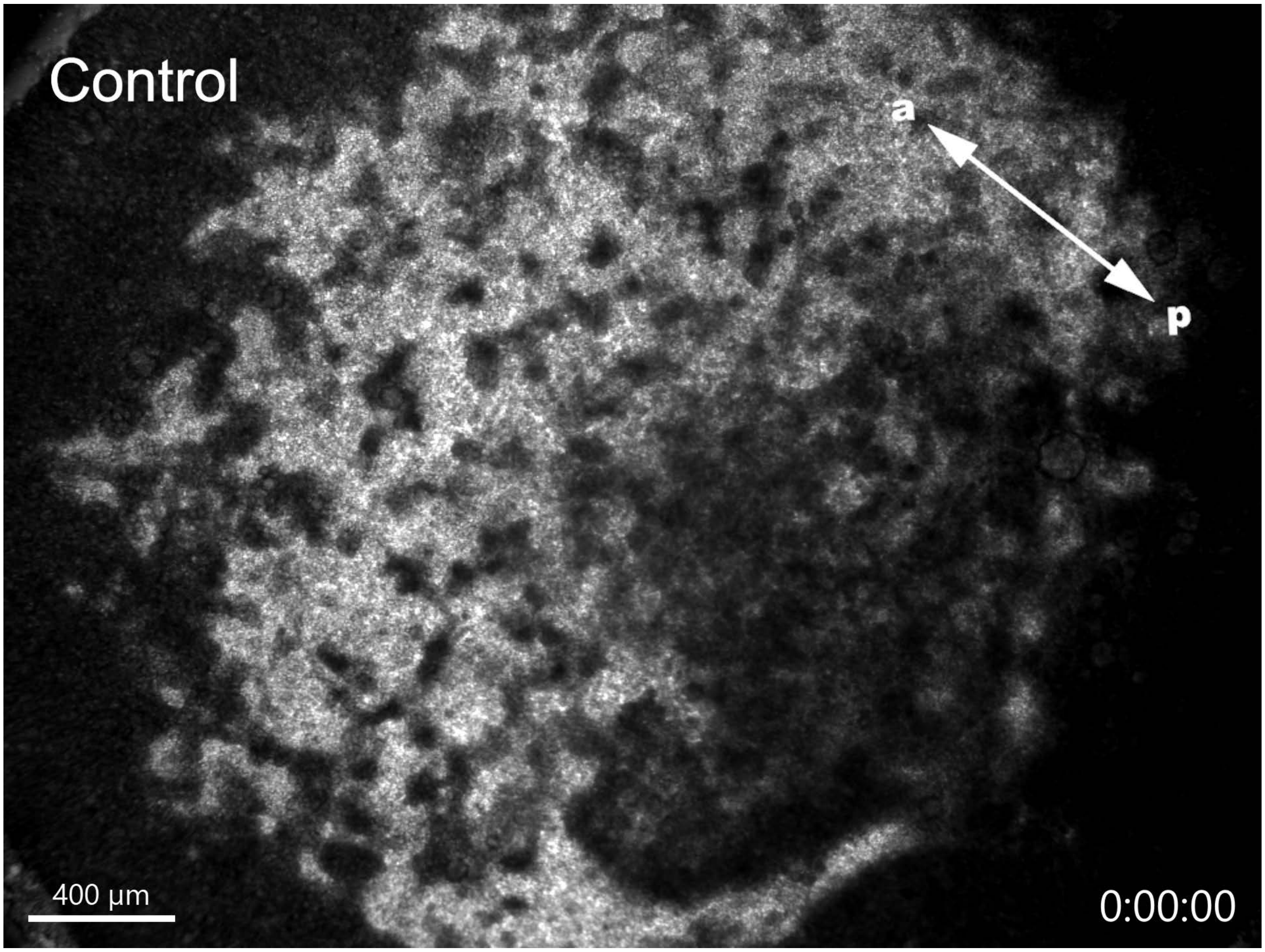


**Movie S1. Brightfield timelapse of pre-streak stage chicken embryo treated with DMSO/control.**
Time-lapse movie taken from pre PS-formation (stage EGK X) to stage HH3 where primary PS was formed. Anteroposterior axis indicated by two-head arrows; location of PS formation is indicated once it starts to form. Live imaging performed at 4x objective lens on a widefield epifluorescent microscope until HH3. Related to Fig. 1. Scale bars, 400μm.
