## supplemental movie 2 for "Differential Sensitivity of Midline Patterning to Mitosis during and after Primitive Streak Extension"

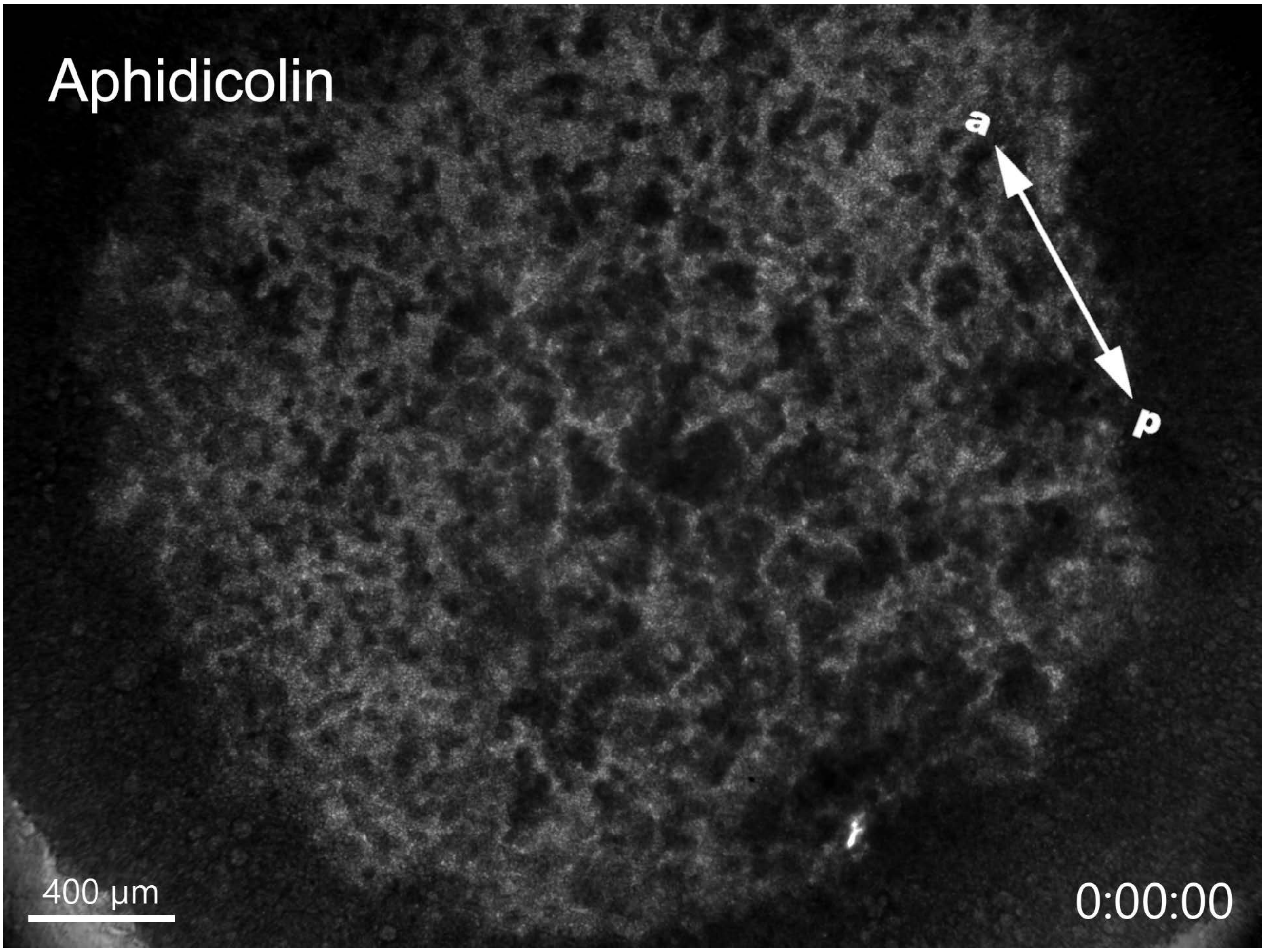


**Movie S2. Brightfield timelapse of pre-streak stage chicken embryo treated with Aphidicolin.** Time-lapse movie of pre PS-formation (stage EGK X) embryo that was treated with Aphidicolin. Anteroposterior axis indicated by two-head arrows. Live imaging performed at 4x objective lens on a widefield epifluorescent microscope for 12hr. Related to Fig. 1. Scale bars, 400µm.
