## supplemental movie 3 for "Differential Sensitivity of Midline Patterning to Mitosis during and after Primitive Streak Extension"

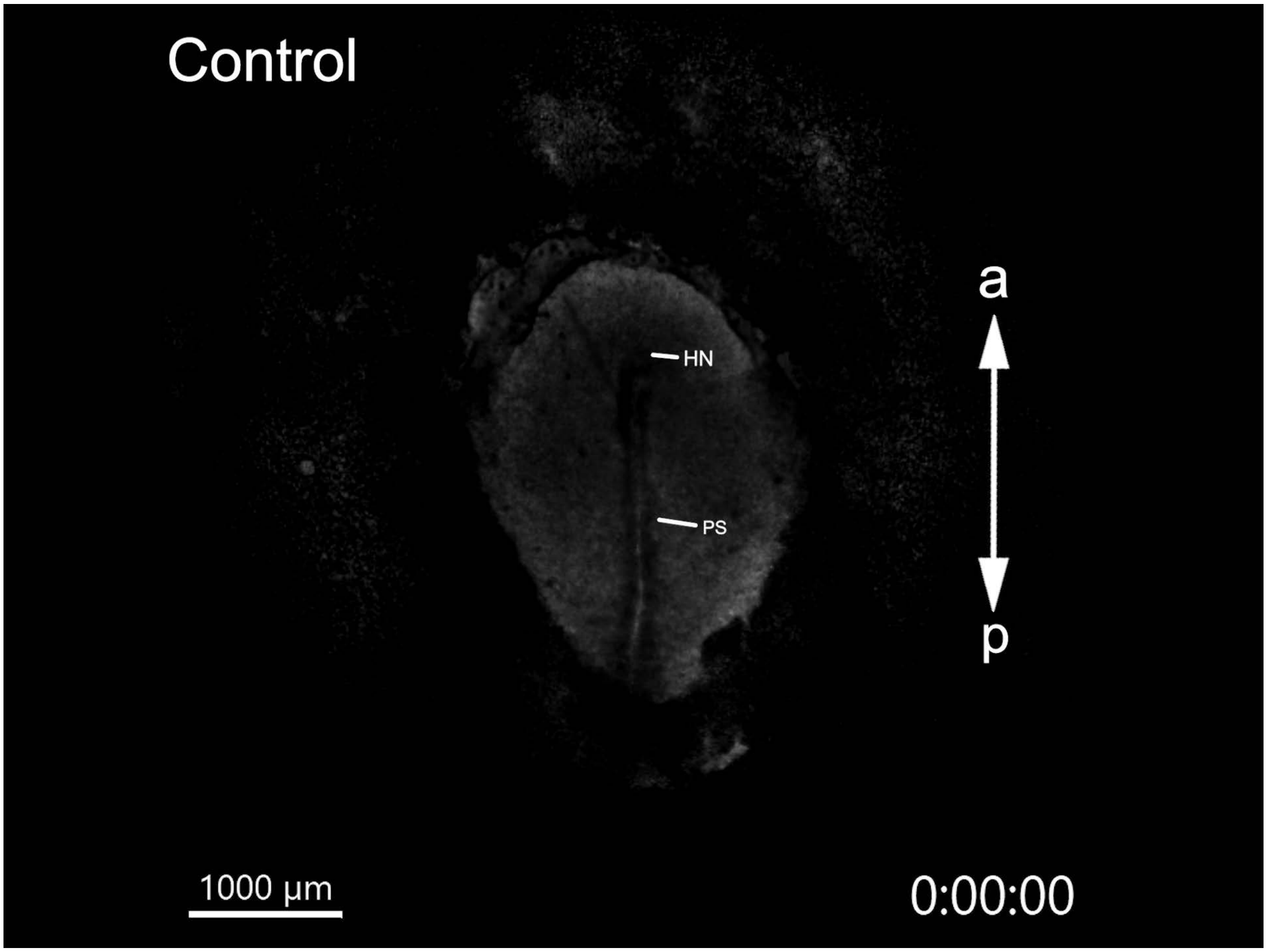


**Movie S3. Brightfield timelapse of post PS-extension stage (HH4) chicken embryo in DMSO/control.**
Time-lapse movie taken from post PS-extension (stage EGK X) to stage HH8+ where NC forms and HN/PS regresses. Anteroposterior axis indicated by two-head arrows; location of NC, HN, and PS are indicated by white bars. Live imaging performed at 2x objective lens on a widefield epifluorescent microscope for 15hr. Related to Fig. 1, 5, Figure S2, and Table1. Scale bars, 1000µm.
